# Socioeconomics and biogeography jointly drive geographic biases in our knowledge of plant traits: a global assessment of the Raunkiærian shortfall in plants

**DOI:** 10.1101/2022.09.26.509556

**Authors:** Brian Maitner, Rachael Gallagher, Jens-Christian Svenning, Melanie Tietje, Elizabeth H. Wenk, Wolf L. Eiserhardt

**Affiliations:** University at Buffalo, Department of Geography, 125a Wilkeson Quadrangle, 14261 Buffalo, NY, United States; Hawkesbury Institute for the Environment, Western Sydney University, Locked Bag 1797, Penrith, NSW 2751, Australia; Center for Biodiversity Dynamics in a Changing World (BIOCHANGE); Section for Ecoinformatics and Biodiversity, Department of Biology, Aarhus University, Ny Munkegade 114, DK-8000 Aarhus C, Denmark; Evolution & Ecology Research Centre, School of Biological, Earth, and Environmental Sciences, UNSW Sydney, Sydney, Australia; Royal Botanic Gardens, Kew, Richmond, Surrey, UK

**Keywords:** Functional Traits, Spatial biases, Data availability, Raunkiæran shortfall, Raunkiærian shortfall, Raunkiaerian shortfall

## Abstract

The traits of plants determine how they interact with each other and their environment, constituting key knowledge for diverse fields. The lack of comprehensive knowledge of plant traits (the “Raunkiærian shortfall”) poses a major, cross-disciplinary, barrier to scientific advancement. Spatial biases in trait coverage may also lead to erroneous conclusions affecting ecosystem management and conservation planning. Thus, there is an urgent need to assess the spatial completeness of plant trait data, understand drivers of geographic biases, and to identify solutions for filling regional gaps. Here, we leverage a comprehensive set of regional species checklists for vascular plants and trait data for 2,027 traits and 128,929 plant species from the TRY database to assess trait data completeness across the globe. We show that trait data availability in TRY is associated with socioeconomic and biological factors influencing sampling likelihood: trait completeness was positively associated with mean species range size, research expenditure, and human population density and negatively associated with endemism and vascular plant species richness. Integration of a second, regional trait database (AusTraits) more than doubled trait completeness for the continent covered, indicating that the creation and integration of regional databases can rapidly expand trait completeness.

**Plain Language Summary:** The traits of plants determine how they interact with each other and their environment. Our knowledge of plant traits is incomplete, limiting scientific advancement as well as our ability to manage ecosystems and plan conservation actions. We show that there are large biases in trait data availability which are associated with both biological factors (range size, endemism, species richness) and socioeconomic factors (research expenditure, human population density). We also show how regionally-focused efforts can help rapidly expand trait data availability.

## Introduction

The traits (measurable attributes) of organisms determine how they interact with their biotic and abiotic environment. Traits allow us to understand both how individuals and the communities they form will respond to environmental change and how these changes will impact ecosystem services and processes (Lavorel & Garnier, 2002). Plants constitute the vast majority of life on Earth (~82% by biomass; Bar-On *et al*., 2018), and their traits are the predominant drivers of terrestrial ecosystem functioning (Migliavacca *et al*., 2021; Fricke *et al*., 2022). Thus, to a first-order approximation, understanding the traits of plants means understanding terrestrial ecosystems.

There remains a sustained interest in both trait-based ecology (e.g., Lavorel & Garnier, 2002; McGill *et al*., 2006; Violle *et al*., 2007; Mouillot *et al*., 2021) and Open Science (Cheruvelil & Soranno, 2018; Gallagher *et al*., 2020b; Geange *et al*., 2021), both of which have contributed to the creation and sharing of large compilations of plant traits constituting millions of observations (e.g. Kattge *et al*., 2011; Maitner *et al*., 2017; Sauquet *et al*., 2017; Falster *et al*., 2021). However, despite this growing wealth of data, our knowledge of plant traits remains far from complete (the ‘Raunkiaeran shortfall’; Hortal *et al*., 2015).

Recent work by Cornwell et al. (2019) examined the coverage of a range of attributes, including traits, in the global flora with a focus on assessing the completeness (presence/absence) of information using The Plant List as a taxonomic backbone. Here, we expand on this work by mapping trait completeness globally, using the geographic and taxonomic information in the recently completed World Checklist of Vascular Plants (WCVP; Govaerts *et al*., 2021) and trait information from the widely-used TRY database (the most commonly used global plant trait database; Kattge *et al*., 2011). We test hypothesized drivers of variation in plant trait data availability across the globe, including factors related to 1) wealth and educational expenditure, 2) region size and accessibility; and 3) biogeography. We compare spatial patterns of trait data completeness with those of phylogenetic and distributional data completeness (Rudbeck *et al*., 2022). Finally, we identify solutions for filling significant regional gaps in trait data completeness using Open Science approaches, highlighting the model developed for the AusTraits database (Falster *et al*., 2021).

## Methods and Materials

### Data Sources

Species taxonomic and distributional data were taken from the World Checklist of Vascular Plants (WCVP), a global database of all known vascular plant species. The WCVP maps plant ranges to the “botanical countries’’ of the World Geographical Scheme for Recording Plant Distributions (formerly known as Taxonomic Databases Working Group, TDWG; Brummitt *et al*., 2001; Banki *et al*., 2019). Following Rudbeck et al. (2022), we filtered the WCVP to include only accepted species occurring within botanical countries in which they were both native and extant. We considered the set of accepted names from the WCVP as our global species pool and the set of species assigned to a botanical country as the species pool for that region.

We obtained all publicly available plant trait data from the TRY database (Kattge *et al*., 2011) on 10 March, 2022. Names for the taxa within the TRY database were standardized to the WCVP taxonomy using the Taxonomic Name Resolutions Service (TNRS; Boyle *et al*., 2013; Maitner & Boyle, 2022) with the WCVP as the source. We focused our primary analyses on traits that could be measured on any vascular plant and omitted traits which were not attributes of the individual (e.g., population density, native status, human use). In order to exclude traits with few observations but still include a large variety of traits, we excluded traits for which less than one percent of vascular plant species had data globally (n = 55 traits included). This primary set of trait data was combined with the WCVP checklist to calculate and map the percent completeness of each trait within each botanical country. In additional analyses, we also assessed completeness of georeferenced trait data.

Following Rudbeck et al. (2022), we focused on three classes of predictor variables expected to be associated with trait data availability, including those related to: 1) funding; 2) accessibility; and 3) biogeography. Funding-related predictors included total Gross Domestic Product (GDP; Giuliani & Peduzzi, 2011), per capita GDP (Giuliani & Peduzzi, 2011), research expenditure (World Bank World Development Indicators, 2017), and education expenditure (World Bank World Development Indicators, 2016). Accessibility-related predictors were area, road density (Meijer *et al*., 2018), security (Institute for Economics and Peace, 2019), population size (Center for International Earth Science Information Network (CIESIN), 2018), and population density (Center for International Earth Science Information Network (CIESIN), 2018). Biogeographic predictors included species richness, mean range size, and endemism and were calculated from the WCVP (Govaerts *et al*., 2021), with endemism calculated following the methods in Gallagher et al. (Gallagher *et al*., 2020a). Species richness was calculated as the number of species within a region and endemism as the number of endemic species divided by species richness. The range size of each species was estimated as the total area of the botanical countries that a species occurred in, with the mean range size of a region being the average of these range sizes across species (Rudbeck *et al*., 2022). All predictor variables were rescaled to a mean of zero and a variance of one.

### Analyses

All analyses were conducted in R version 4.1.2 (R Core Team, 2020). We used generalized, linear, mixed-effects models to test for a relationship between predictor and response variables. The response variable followed a binomial distribution with successes defined as species with data and failures as species without data. Predictor variables included: GDP (sum), GDP (per capita), research expenditure, education expenditure, area, road density, population, population density, security, species richness, mean species range area, and endemism. Trait name was included as a random effect. Preliminary analyses indicated significant spatial autocorrelation in the response variable (Moran’s I = 0.41, p < 2.2e-16), and so we also included a Matern random effect to account for spatial autocorrelation (Stein, 1999). Models were built using the function `fitme` in the R package *“spaMM”* (Rousset & Ferdy, 2014).

Trait data increase whether the traits of a species are measured within a focal botanical country or another region in which it occurs, weakening relationships between predictor and response variables. To account for this confounding effect of shared species, we repeated our analyses on the georeferenced subset of trait data. We excluded any trait data which could not be georeferenced to a botanical country on the basis of either a declared political division name or latitude and longitude, or where the coordinates disagreed with the declared political division. For this analysis, trait data only counted toward the completeness of the botanical country in which they were recorded. Coordinates were parsed using the R package *“parzer”* (Chamberlain & Sagouis, 2021) and standardization of unmatched botanical country names was attempted using the R package *“GNRS”* (Boyle & Maitner, 2021). As above, models were built using the function `fitme` with a binomial response (species with data/without data) and the same predictor variables and random effects.

We tested for correlations between completeness of trait, phylogenetic, and distributional data across botanical countries (Pearson correlation) by comparing our results with those of Rudbeck et al. (2022). For these analyses, trait completeness was calculated as the mean completeness of all traits with data for at least 1% of species globally (n = 55) and our species pool was limited to Spermatophytes for consistency with Rudbeck et al. Phylogenetic and distributional completeness was taken from Rudbeck et al. (2022), with phylogenetic completeness calculated as the fraction of species × genetic marker combinations with data and distributional completeness calculated as the fraction of species in a region with coordinate data in the BIEN database (Enquist *et al*., 2009; Maitner *et al*., 2017) for that region.

Finally, we assessed the degree to which a regionally-focused trait gathering initiative, AusTraits (Falster *et al*., 2021), could improve trait data completeness relative to TRY alone. We downloaded AusTraits version 3.0.2 using the *“austraits”* R package, standardized trait names to match those in TRY, and standardized taxonomy to the WCVP using the *“TNRS”* R package (Boyle *et al*., 2013; Maitner & Boyle, 2022). We retained trait data for our 55 focal traits, and combined it with that already present in TRY to assess the magnitude of improvement in trait completeness across Australia when integrating the two databases (Feng *et al*., 2022).

## Results

**Figure 1.**
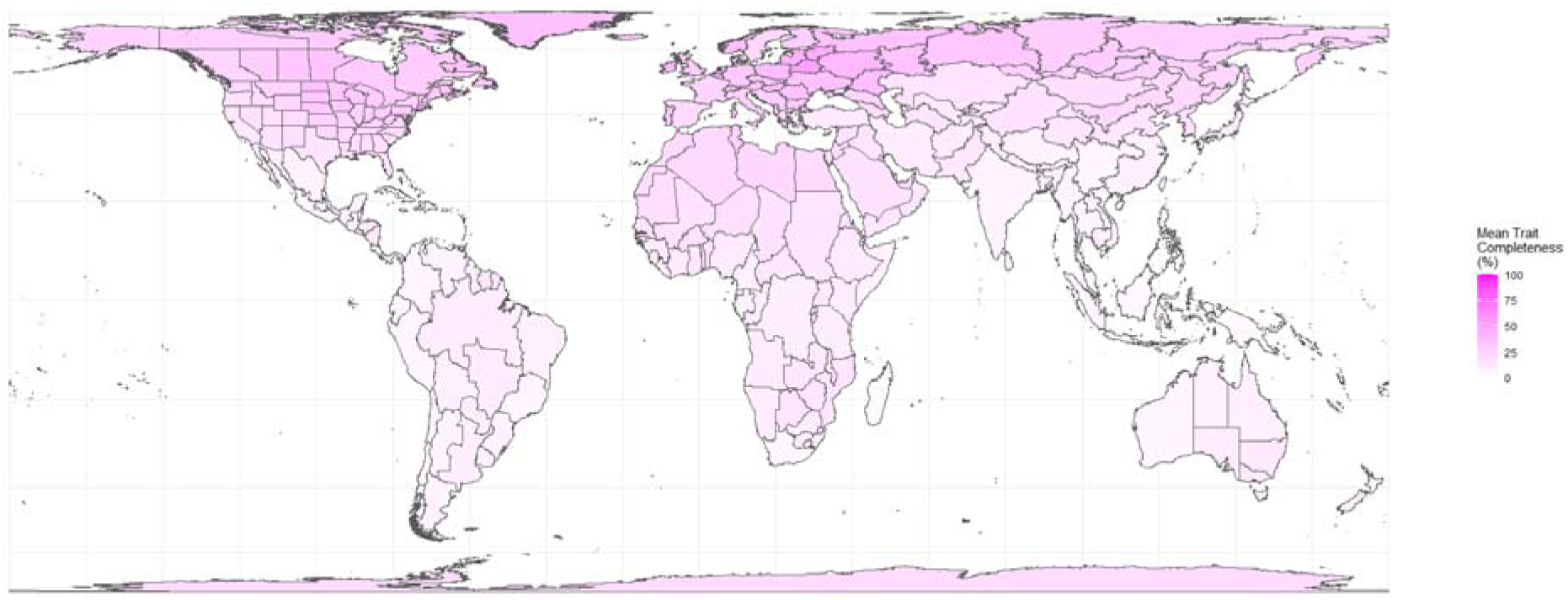
Mean Trait Completeness in the TRY database based on the World Checklist of Vascular Plants. Trait completeness was calculated as the number of species with trait information for a given trait divided by the total number of species in the botanical country. Mean completeness was calculated across the set of 55 focal traits that could be applied to all vascular plants and which contained data for at least one percent of species globally.

The set of public trait data we received from TRY contained a total of 2,027 unique traits, with the most complete being plant growth form (31.9% of species). Most traits had poor coverage (Figure SI 1), with a mean global completeness across all traits of 0.20% and a median of 0.0049%. Surprisingly, some traits calculated from multiple measurements had higher coverage than their components (e.g., SLA was available for 14,127 species, leaf dry mass for 6,892). After excluding traits with data for less than one percent of species, our focal dataset included 55 traits ranging between 1.05% and 31.9 % global coverage (mean = 3.55%, median = 1.87%). For a single trait, within a single botanical country, trait completeness varied widely, ranging between 0 and 100% (mean = 18.8%, median = 12.2%), with mean completeness across traits ranging between 5.9 and 69.8% across countries (mean = 18.8%, median = 15.3%, Figure 1). Correlations in completeness across botanical countries were positive and significant for the majority of focal traits, with the exception of a few leaf morphological traits (width, length, margin type, venation type) and species habitat characterization: vegetation type (Figure SI 2).

We found that the effects of five of our predictor variables had 95% confidence intervals that excluded zero (Table SI 1). Trait completeness was positively associated with mean species range size (0.40, 95% CI [0.36, 0.44]), research expenditure (0.06, 95% CI [0.03,0.09]), and human population density (0.02, 95% CI [0.001, 0.04]). Trait completeness was negatively associated with endemism (−0.13, 95% CI [−0.15, −0.10,]) and vascular plant species richness (−0.09, 95% CI [−0.12, −0.06]).

Of the 2,027 unique trait names in the TRY dataset, 1547 (76%) contained at least one georeferenced trait observation, amounting to a total of 613,492 species × trait × botanical country combinations. Coverage of georeferenced data was considerably worse than for trait data in general, with maximum trait coverage of 5.46% (for plant growth form), and a mean coverage of 0.07%. After excluding traits with data for less than one percent of species, our georeferenced dataset included 29 traits ranging between 1.04% and 5.46% coverage (mean = 1.8%, median = 1.6%). There was a weak, positive correlation between georeferenced trait completeness and general trait completeness (r = 0.15, p = 0.005, Figure SI 4). With the exception of mean species range size (−0.22, 95% CI [−0.48, 0.30]), all of the hypothesized predictors had confidence intervals that excluded zero (Table SI 2). Georeferenced trait completeness was positively related to research expenditure (1.32, 95% CI [1.25, 1.60]), species richness (0.97, 95% CI [1.00, 1.29]), area (0.90, 95% CI [0.87, 1.01]), GDP(sum) (0.40, 95% CI [0.38, 0.52]), security (0.27, 95% CI [0.16, 0.43]), GDP (per capita) (0.22, 95% CI [0.14, 0.63]), education expenditure (0.15, 95% CI [0.05, 0.25]), and population size (0.02, 95% CI [0.004, 0.05]). Georeferenced trait completeness was negatively related to road density (−0.26, 95% CI [−0.40, −0.08]), population density (−0.24, 95% CI [−0.55, −0.13]), and endemism (−0.09, 95% CI [−0.24, −0.06])

At the level of botanical countries, trait completeness showed a strong, positive correlation with phylogenetic completeness (r = 0.81, p < 0.01; Figure 2, Figure 2 A) but was not significantly correlated with distributional completeness (r = 0.05, p = 0.34, Figure 2 C). Distributional data tended to be the most complete across the globe (mean = 47.12%, median = 51.64%), although for some regions trait data were more complete (e.g., Selvagens Islands, Ascension Island, Wake Island, and much of Russia; Figure 2, C and D), however, phylogenetic data were always less complete than trait data (Figure 2 B).

**Figure 2.**
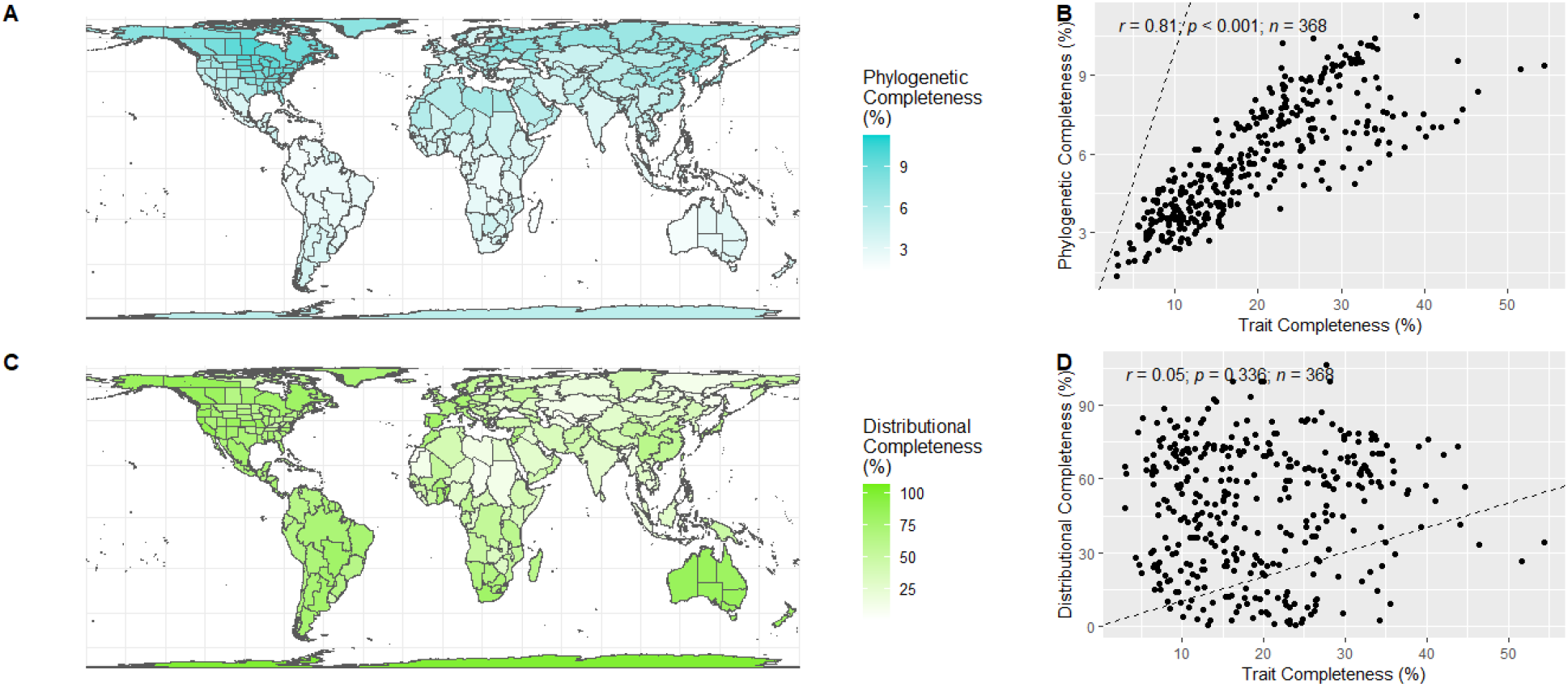
Correlations among biodiversity shortfalls across botanical countries. A. Phylogenetic completeness refers to the percent of 128 commonly sequenced genetic regions available for species within a botanical country. C. Distributional completeness refers to the availability of any georeferenced point occurrence data in the BIEN database for the species within a botanical country. Distributional and genetic completeness were calculated following Rudbeck et al. (2022). B and D: Dashed lines indicate the 1:1 line and points. For B and D, ferns were excluded from calculations of trait completeness to insure comparability with Rudbeck et al. (2022).

For Australian vascular plants, the TRY database contained information on 71,324 species x trait combinations across 9,722 species and all 55 focal traits. After filtering the AusTraits database to only include our focal traits, it contained a total 171,219 species x trait combinations across 20,499 species and 35 of our focal traits. When integrated, the two databases contained 200,891 species x trait combinations across 21,264 species (99.5% of Australian vascular plants, per the WCVP). This increase in data resulted in a more than doubling of mean trait completeness across all of the botanical countries (i.e., states) within Australia (Figure 3).

**Figure 3.**
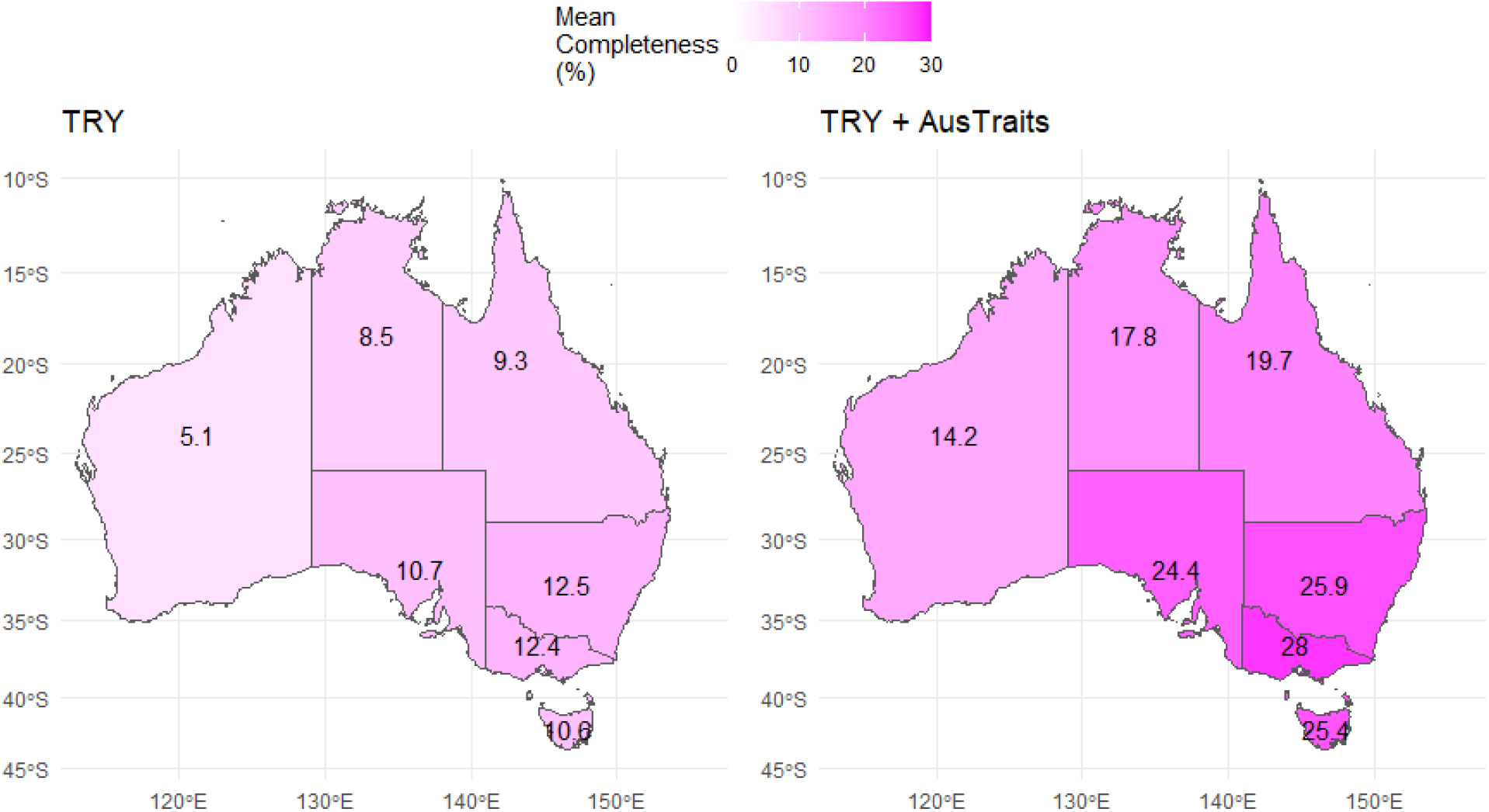
Database integration improves trait completeness. Trait completeness was calculated as the number of species with trait information for a given trait divided by the total number of species in the botanical country. Mean completeness was calculated across the set of 55 focal traits that could be applied to all vascular plants and which contained data for at least one percent of species globally. The numbers within each botanical country are the mean trait completeness (percent).

## Discussion

Despite the massive amount of trait data collated to date, we are a long way from fully capturing some of the simplest traits for the majority of plant species across the world’s botanical countries. All 55 focal traits examined here were below the 40% coverage threshold noted by Penone et al. (2014) as being needed to impute missing traits with confidence, suggesting that imputation is not yet an option at a global level (but may be of use within certain regions). Our mapping shows that what information we do have is spatially biased, with trait coverage being higher in the Global North. Completeness is generally consistent across traits, showing that we have a general lack of trait data in the Global South, as also noted by Cornwell et al. (2019), rather than simply a different set of traits being measured. However, we note that 19% of traits show a negative correlation with mean focal trait completeness (Figure SI 3), but most of these were below our 1% threshold for inclusion. The geographic biases we observe are similar to those observed for phylogenetic data by Rudbeck et al. (2022), with the factors we identified as driving trait data completeness (range size, endemism, species richness, research expenditure, and population density) being the same factors they identified as driving phylogenetic data completeness, suggesting similar mechanisms underlie the acquisition of both data types. Unfortunately, this correspondence between phylogenetic and trait data completeness likely means that regions with low trait completeness have relatively less to gain trait data via phylogenetic trait imputation.

Comparing our main analysis, which included traits measured anywhere, with our analysis focusing only on traits known to have been measured within particular botanical countries reveals socioeconomic and biological factors contributing to these global biases. By removing the confounding influence of shared trait data, the georeferenced data provide a better picture of the drivers of trait data availability at a botanical country level, and show that trait data completeness is positively associated with national wealth and spending, with wealthier nations that spend more on research and education typically having better coverage. We note that in our main analysis (Figure 1), some countries with high levels of wealth and spending appear relatively data poor (e.g., Australia and New Zealand). However, when we combine AusTraits with TRY, the level of trait completeness in Australia falls in line with levels seen elsewhere in the Global North (Figure 3), suggesting that a lack of data integration may underlie some perceived knowledge gaps. In addition to relatively high rates of funding, countries in the temperate and polar regions of the Northern Hemisphere also have relatively low species diversity and large species ranges, allowing them to share data across boundaries. In contrast, many tropical countries have low rates of funding, high endemism, and high species diversity, factors which work against trait completeness. Although the focus of this study was botanical countries, the bias we observe will extend to other geographic classifications (e.g., biogeographic realms, biomes, ecosystems; Olson *et al*., 2001) such that our knowledge of those occurring predominantly in the Global South will tend to be relatively poor.

The analyses presented here focus on the most widely-used global plant trait database, TRY (Kattge *et al*., 2011, 2020). However, numerous other plant trait databases and datasets exist, some of which will not have been incorporated in TRY yet, and others which may not be able to be incorporated due to licensing issues. However, by integrating TRY with one open access resource (AusTraits), we were able to more than double trait data completeness for all Australian states (Figure 3). This supports recent work by Feng et al. (2022) showing that database integration can lead to rapid gains in available data. The AusTraits model demonstrates how the creation and integration of regional databases can rapidly expand trait completeness. With a more limited geographic scope, AusTraits could target its data collection, allowing it to achieve similar completeness to the northern hemisphere despite high endemism and a low population density. This includes personally interacting with researchers to create a sense of community, developing a workflow where a database manager leads the input of datasets reducing contributor effort, and contacting researchers and repositories with known large datasets. In particular, AusTraits makes use of expertise held within the systematics community, mining data embedded in taxonomic descriptions. This approach provides depth of coverage for key traits such as growth from, leaf dimensions, and plant height aiding in the filling of regional gaps for modeling and conservation management. Due to socioeconomic limitations, regional efforts in some areas (particularly in the Global South) may only be feasible with South-South and South-North collaborative efforts. Data source integration is currently hindered by variation in data structure, data availability, taxonomy, and even trait names (Gallagher *et al*., 2020b), so new, regional efforts would benefit from adopting existing tools, taxonomies, and data structures (e.g., Boyle *et al*., 2013; Schneider *et al*., 2019; Falster *et al*., 2021).

In this study we quantified trait data completeness relative to a subset of traits provided in the TRY database, but there are an infinite number of possible traits: measurements can be taken at any point, organ or developmental stage on an individual, at multiple levels of organization, and combined in any way (e.g., dry leaf mass per leaf area, above ground biomass divided by belowground biomass). A lack of standard trait definitions makes the integration of different databases challenging. Thankfully, plant traits are often strongly correlated (Westoby *et al*., 2002; Wright *et al*., 2004; Díaz *et al*., 2016; Zeballos *et al*., 2017), and even sparse trait coverage may allow us to say something about the overall phenotypes of species (Mouillot *et al*., 2021), particularly when combined with phylogenetic information (Penone *et al*., 2014) or geographic information (Sandel *et al*., 2021). Imputation methods based on trait and phylogenetic correlations also provide estimates of uncertainty which can be used in sampling prioritization. Thus, global collection efforts focused on key traits representing known trait spectra (e.g., Westoby *et al*., 2002; Wright *et al*., 2004; Díaz *et al*., 2016; Zeballos *et al*., 2017), particularly in taxa or regions of high uncertainty, may be a reasonable path forward.

While plant traits are critically important across disciplines and are urgently needed to allow us to predict responses to global change, the current state of our knowledge is both incomplete and geographically biased. Given the current state of available data, large scale (e.g., country, biome, global) analyses of plant traits must be interpreted with caution and should attempt to quantify uncertainty caused by these massive data gaps. Moving forward, researchers can help by publishing their data and metadata openly, including the full set of raw and derived trait data (Keller *et al*., 2022). At larger scales, efforts to mobilize and integrate existing datasets, particularly those focused on particular geographic regions (e.g., Tavşanoğlu & Pausas, 2018; Falster *et al*., 2021) or types of traits (e.g. Iversen *et al*., 2017; LeBauer *et al*., 2018; Guerrero-Ramírez *et al*., 2021), hold promise to rapidly advance the state of our knowledge (Feng *et al*., 2022). However, what is ultimately needed is more data collection and research, particularly in the Global South, which requires funding, infrastructure, and capacity building.

## Acknowledgments

The authors wish to thank Daniel Falster for the development of the AusTraits database. JCS considers this work a contribution to his VILLUM Investigator project “Biodiversity Dynamics in a Changing World” funded by VILLUM FONDEN (grant 16549) and JCS’ Center for Ecological Dynamics in a Novel Biosphere (ECONOVO), funded by Danish National Research Foundation. WLE’s contribution was supported by VILLUM FONDEN (grant 0025354).

## Author Contributions

BSM and WLE conceived the initial idea. BSM, WLE, RG, J-CS, and MT planned and designed the research. RG, J-CS, MT, EHW, and WLE provided data. BSM conducted analyses with feedback from all authors. BSM lead the writing with contributions from all authors.

## Data and Code Availability

The data and code used in this manuscript are available at: https://github.com/bmaitner/shortfalls

## Supplementary Information

**Figure SI 1.**
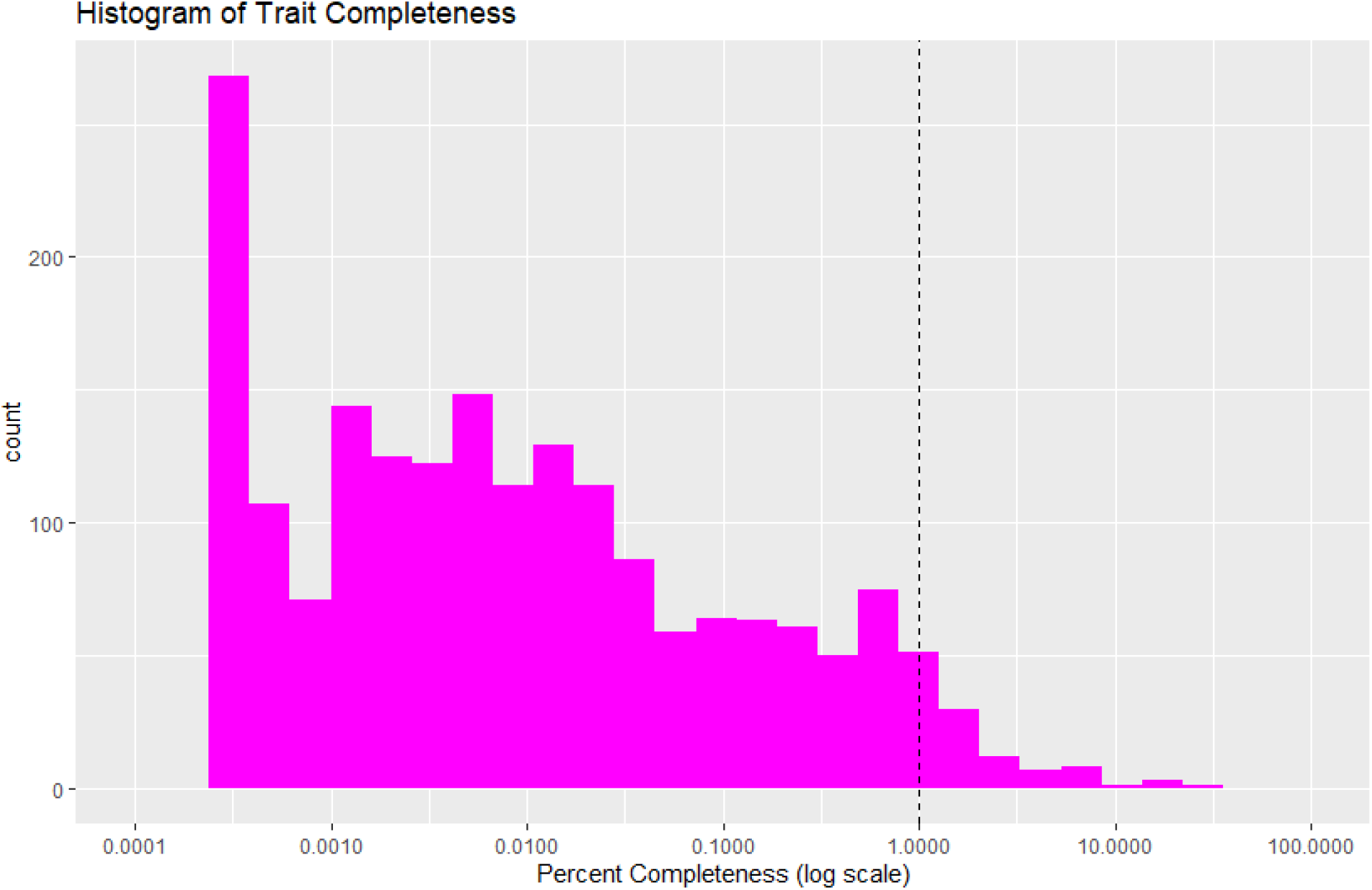
Distribution of trait completeness. Percent trait completeness was calculated as the number of species with data for a given trait divided by the number of species overall, multiplied by 100. The dashed vertical line represents the threshold we set for traits to be included in our analyses.

**Figure SI 2.**
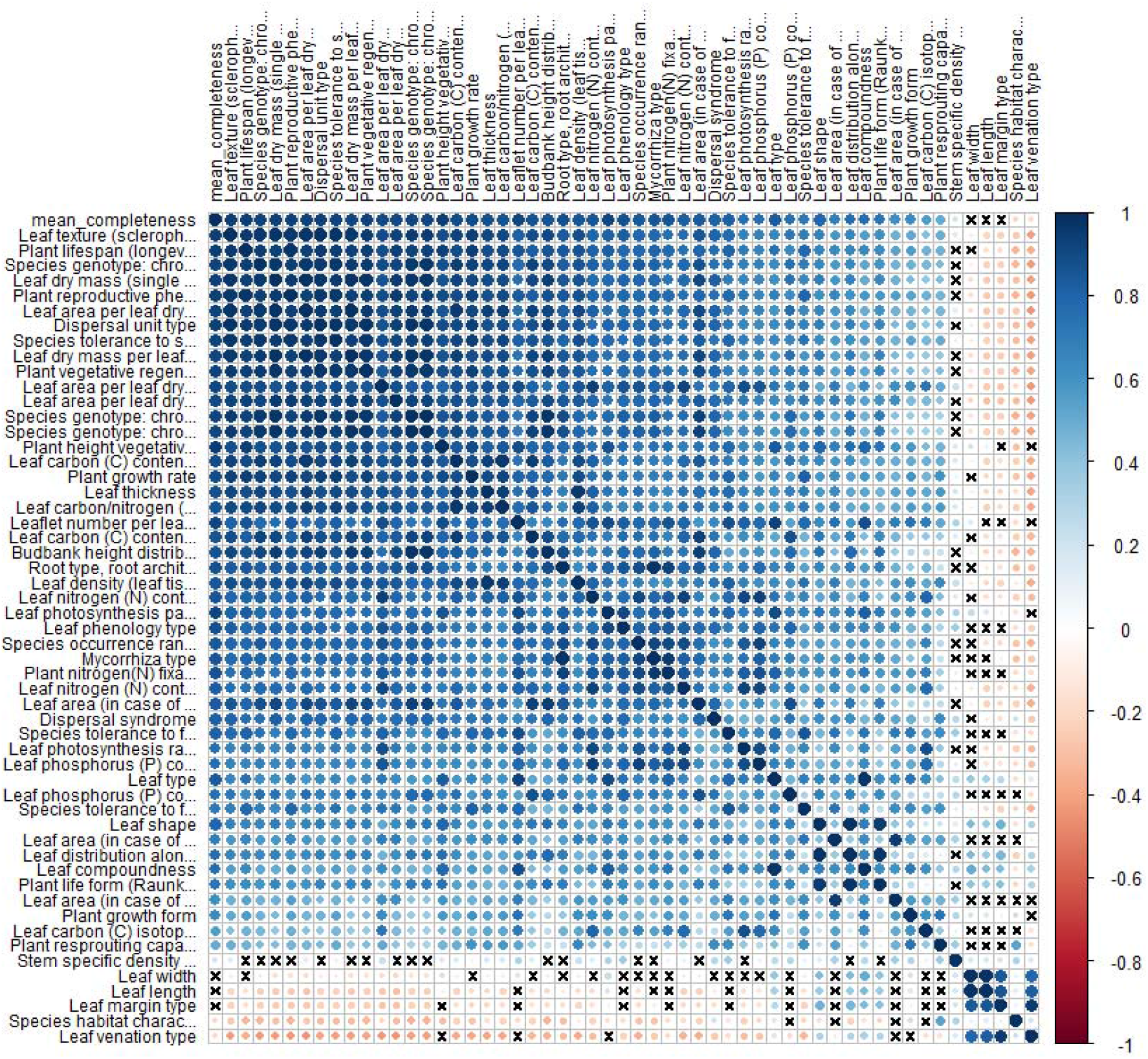
Pearson correlations of trait completeness across botanical countries. Symbol color and size represent the direction and magnitude of correlation. Correlations with an “X” on them are not significant (p > 0.05), all other correlations are significant (p ≤ 0.5).

**Figure SI 3.**
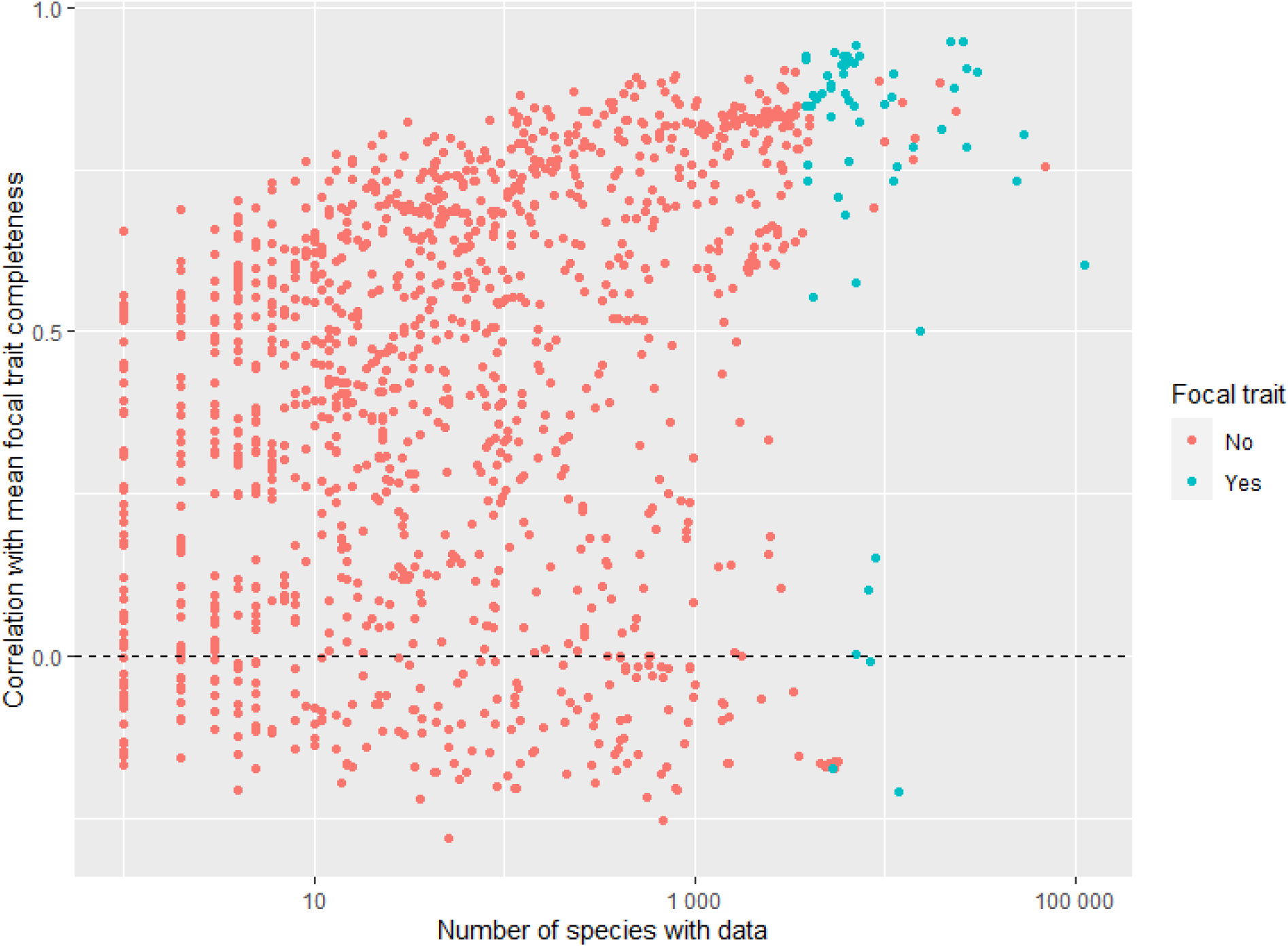
Pearson correlations of trait completeness with mean focal trait completeness. Correlations are calculated across botanical countries. These correlations should be interpreted cautiously, as the underlying completeness estimates are calculated relative to the total set of vascular plant species within a botanical country, regardless of whether the trait can be applied to all species. Focal traits are the set of 55 traits which can be applied to any vascular plant species and for which data are available for at least 1% of species globally.

**Table SI 1.**
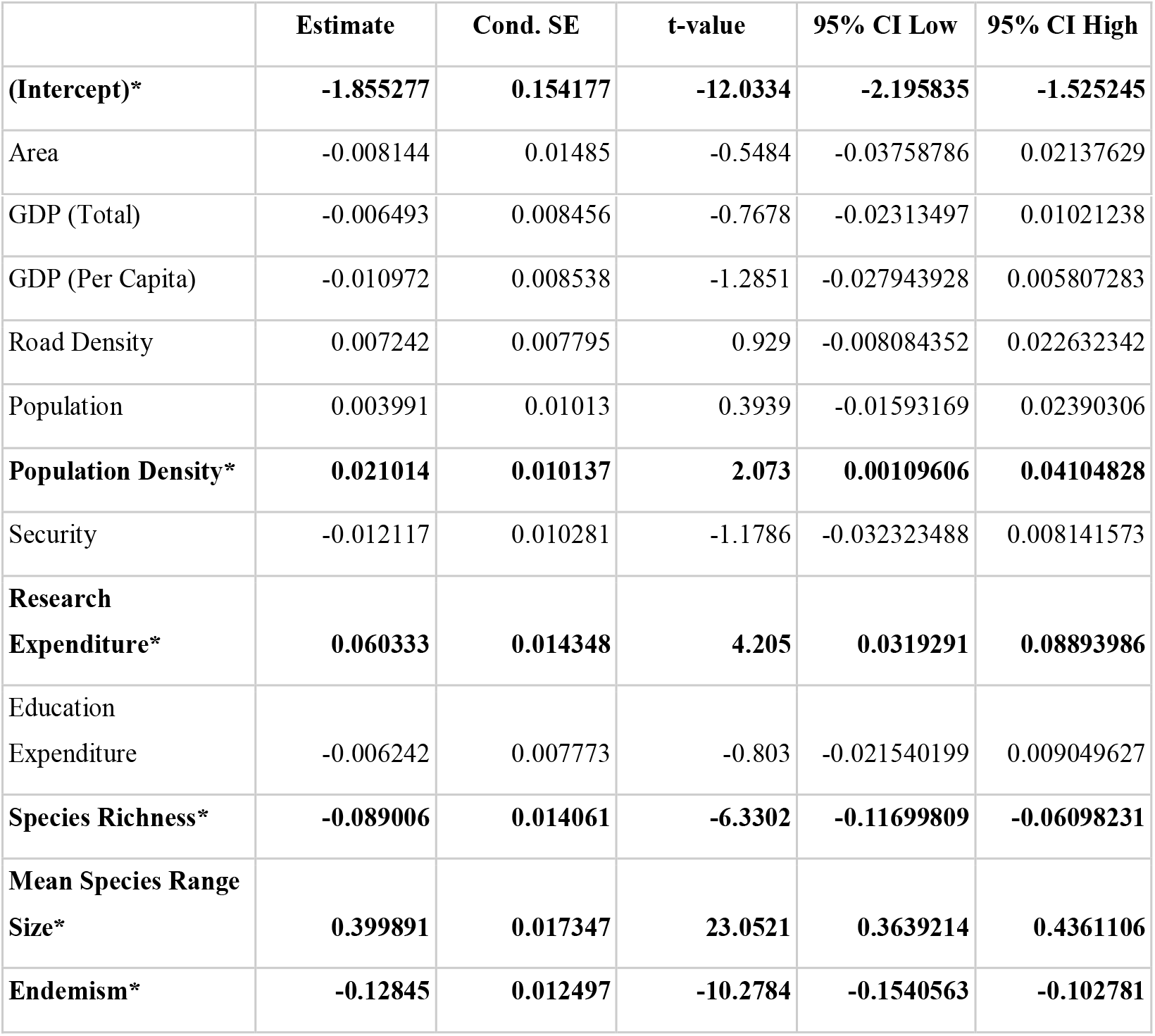
Estimated model coefficients. Model terms in bold and denoted by a * have a 95% CI that does not include zero.

**Figure SI 4.**
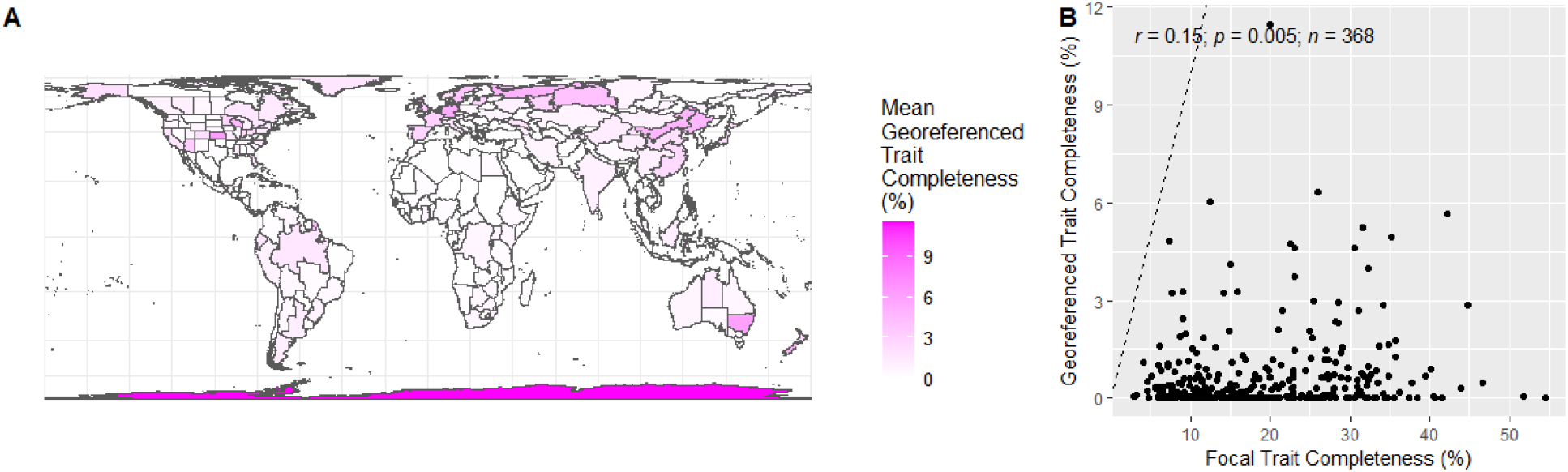
Mean Georeferenced Trait Completeness. A. Georeferenced trait completeness was calculated as the number of species within a botanical country with georeferenced trait information for a given trait divided by the total number of species in the botanical country. Mean georeferenced trait completeness was calculated across the set of 29 focal traits that could be applied to all vascular plants and which contained data for at least one percent of species globally. B.Focal trait completeness was calculated across as in Figure 1. The dashed line indicates the 1:1 line.

**Table SI 2.** Estimated model coefficients for the georeferenced trait model. Model terms in bold and denoted by a * have a 95% CI that does not include zero.

|  | Estimate | Cond. SE | t-value | 95% CI Low | 95% CI High |
| --- | --- | --- | --- | --- | --- |
| **(Intercept)\*** | **-8.22194** | **0.18** | **-45.6787** | **-8.611** | **-8.113** |
| **Area\*** | **0.90282** | **0.1657** | **5.4489** | **0.8749** | **1.0106** |
| **GDP (Total)\*** | **0.39607** | **0.1699** | **2.3313** | **0.3776** | **0.5224** |
| **GDP (Per Capita)\*** | **0.22684** | **0.2655** | **0.8545** | **0.1379** | **0.6281** |
| **Road Density\*** | **-0.25824** | **0.2908** | **-0.8881** | **-0.4031** | **-0.0785** |
| **Population\*** | **0.02202** | **0.1755** | **0.1255** | **0.0048** | **0.0541** |
| **Population Density\*** | **-0.23971** | **0.2692** | **-0.8906** | **-0.551** | **-0.131** |
| **Security\*** | **0.2663** | **0.2084** | **1.2779** | **0.16** | **0.4263** |
| **Research Expenditure\*** | **1.31657** | **0.2315** | **5.686** | **1.246** | **1.596** |
| **Education Expenditure** | **0.15842** | **0.1939** | **0.8172** | **0.0516** | **0.2468** |
| **Species Richness\*** | **0.96996** | **0.2109** | **4.5993** | **0.9979** | **1.2903** |
| Mean Species Range Size | -0.22202 | 0.2152 | -1.0317 | -0.4831 | 0.2969 |
| **Endemism\*** | **-0.08735** | **0.1939** | **-0.4504** | **-0.2369** | **-0.0611** |

## SI Analyses

Since our main analyses excluded traits that could not be evaluated across all vascular plant species, they excluded common traits related to wood, seeds, and flowers. To address this, we carried out a set of secondary analyses testing whether patterns of trait data completeness for these traits were correlated with the patterns observed in our main analysis. To evaluate the completeness of wood traits, we identified woody species using the TRY dataset (Kattge *et al*., 2011). Woody species were inferred based on growth form (containing “tree”, “shrub”, “liana”, or “woody”) or trait name (i.e., containing “tree” or “wood”). Wood traits were identified as those containing “wood” in the trait name. Floral trait completeness was calculated relative to the set of Angiosperm species and floral traits were inferred as those containing “flower” or “inflorescence” in the trait name. Seed trait completeness was calculated relative to the set of Spermatophyte species and seed traits were identified as those containing “seed” in the trait name.

To understand gaps in data availability at the species level, we visualized overlaps in data availability on a species-by-species basis. For this analysis, we assessed whether each species had any trait data (for any of our focal traits), any phylogenetic data (for any of the 128 markers used by Rudbeck et al. (2022)), or any coordinate data. From these data, we generated a Euler plot using the R package *“eulerr”* (Larsson, 2021) to visualize overlap in data coverage.

## SI Results

We inferred 90,048 species as being woody, and identified 175 wood traits, 157 of which had data for at least one species. Coverage across wood traits ranged between one species (0.001% of woody species) to 4.5% of woody species. Eighteen wood traits had at least 1% coverage globally. We inferred 336,712 species as having seeds, and identified 50 seed traits that could be applied to all seed plants. Coverage ranged between 0.002% to 7.12% of seed plants. Only 3 traits had coverage for at least 1% of seed plants. We inferred 335,537 species as having flowers, and identified 36 flower/inflorescence traits. Coverage ranged between 0.0003% (1 species) and 3.75% of species with flowers. Eight traits had coverage for at least one percent of species with flowers. These trait subsets were significantly correlated with our main analysis (seeds: r = 0.87; wood: r = 0.69; flower: r = 0.42; all p << 0.05. Figure SI 4), showing similar geographical patterns (Figures SI 4:5).

**Figure SI 5.**
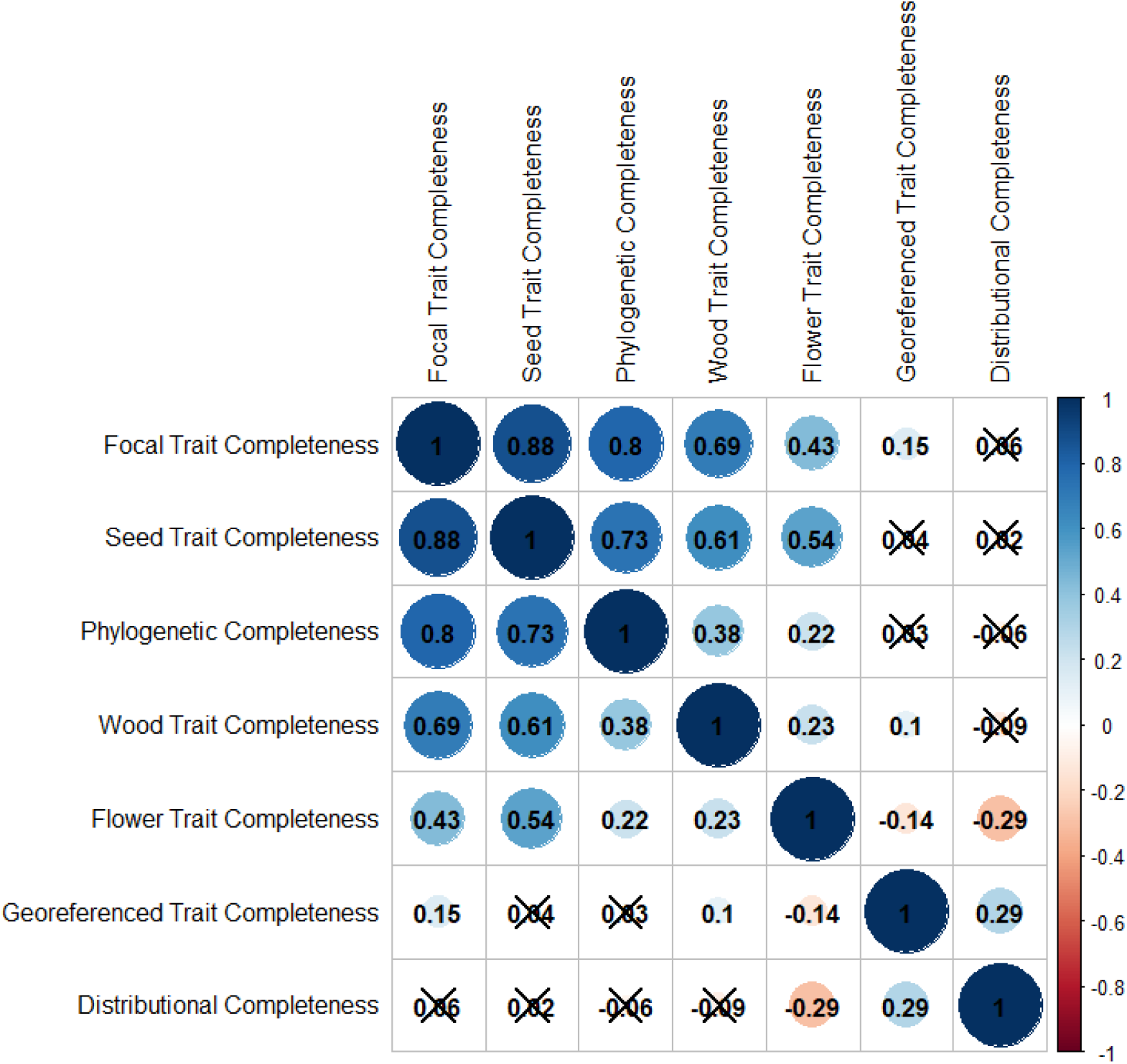
Correlations between completeness for different data types. Values shown are Pearson correlation coefficients calculated between pairs of variables. Symbol color and size represent the direction and magnitude of correlation. Correlations with an “X” on them are not significant (p > 0.05), all other correlations are significant.

**Figure SI 6.**
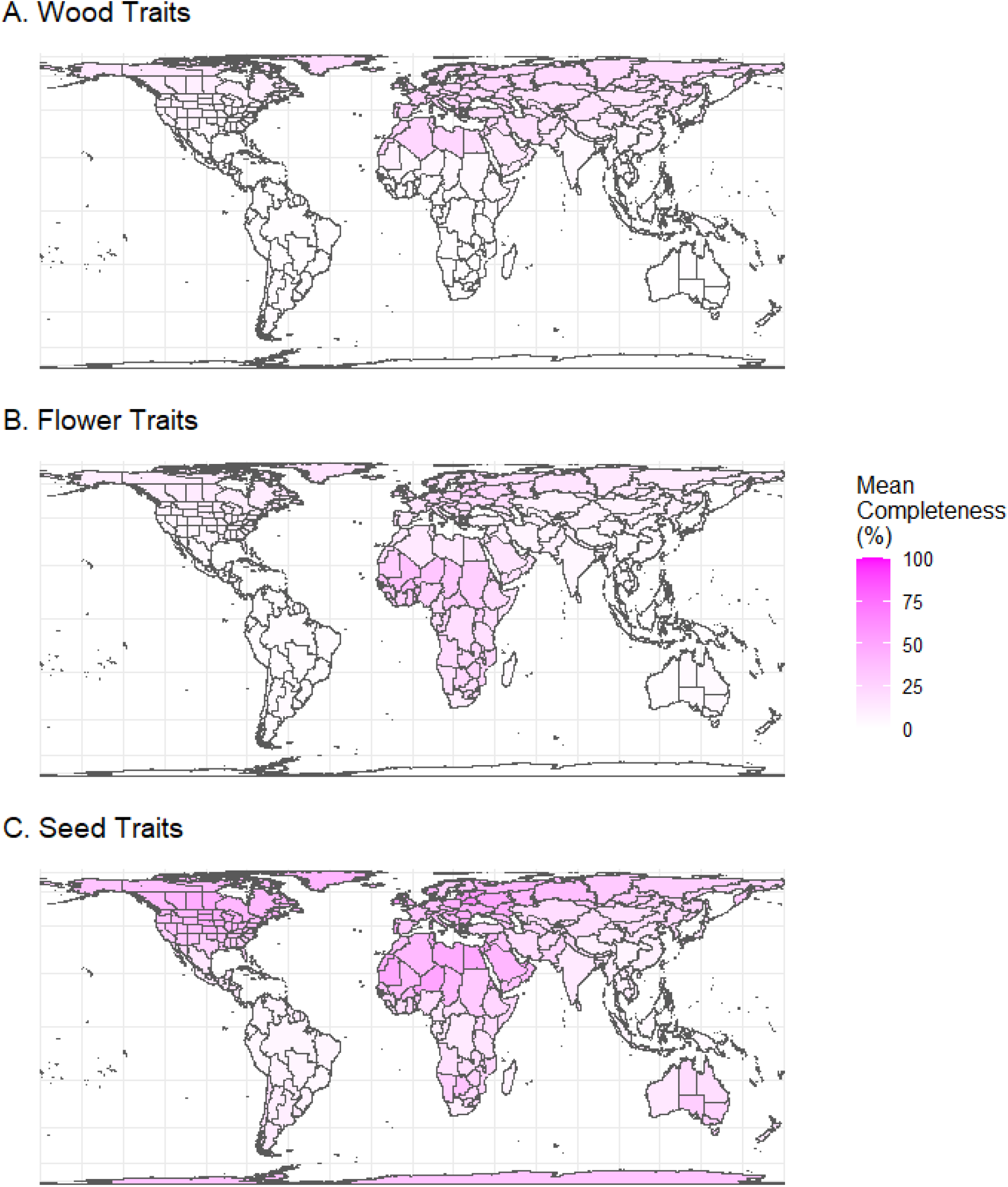
Mean Wood, Flower, and Seed Trait Completeness. Wood trait coverage was calculated as the number of known woody species with trait information for a given trait divided by the total number of woody species in the botanical country. Mean completeness was calculated across the set of 18 wood traits which contained data for at least one percent of species globally. Flower trait coverage was calculated as the number of known Angiosperm species with trait information for a given trait divided by the total number of Angiosperm species in the botanical country. Mean completeness was calculated across the set of 8 flower traits which contained data for at least one percent of species globally. Seed trait coverage was calculated as the number of known Spermatophyte species with trait information for a given trait divided by the total number of Spermatophyte species in the botanical country. Mean completeness was calculated across the set of 3 seed traits which contained data for at least one percent of species globally.

At the level of individual species, there were moderate and significant correlations (Figure SI 6) between trait data availability and both genetic (r = 0.35, p-value < 2.2e-16) and distributional data availability (r = 0.37, p-value < 2.2e-16).

**Figure SI 7.**
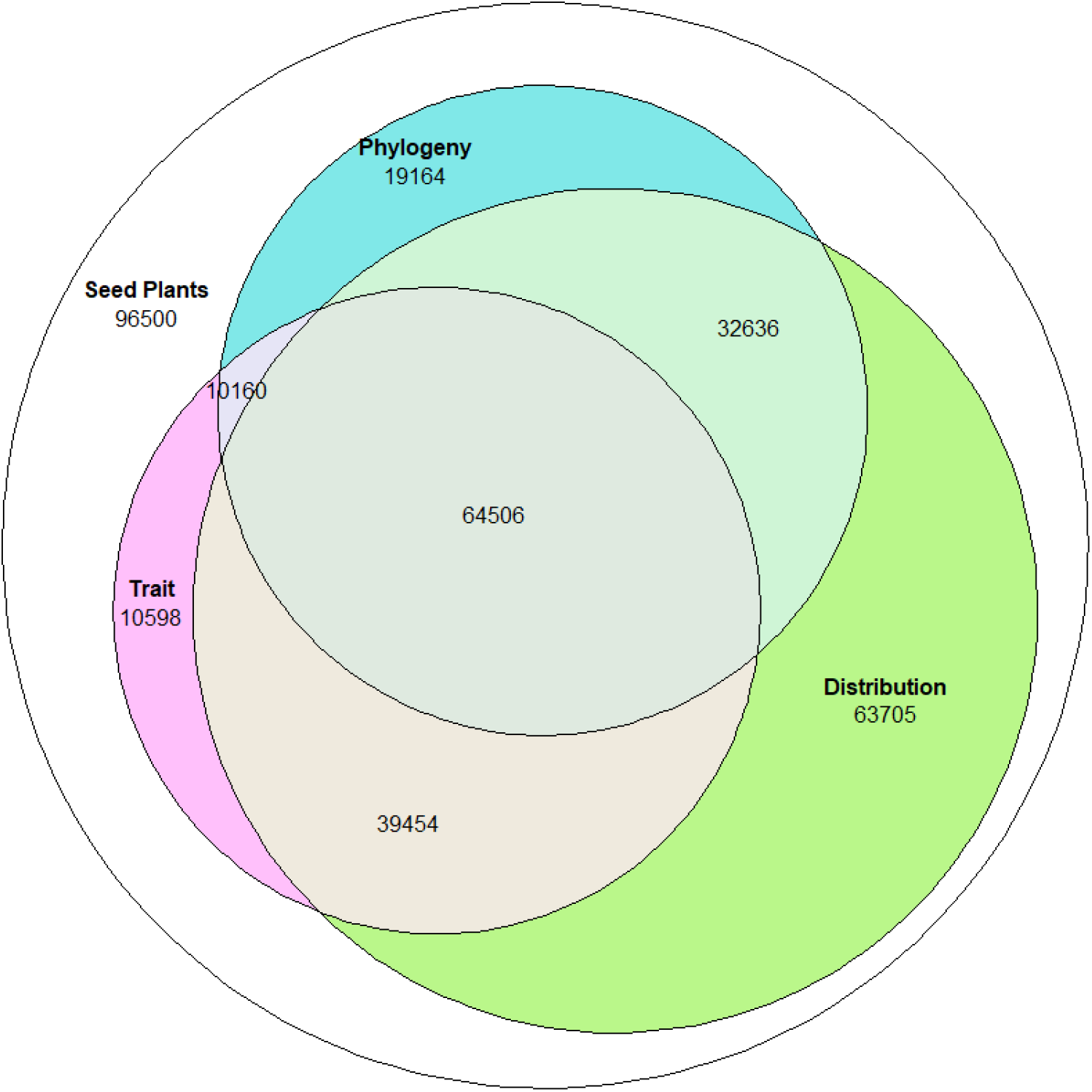
Overlap in biodiversity data availability at the species level. Euler plot with area approximately proportional to overlap. The category “trait” refers to species with any trait data, “distribution” to species with any (validated) coordinate data, “phylogeny” to any species with data in GenBank, and “seed plants” to accepted species present in the WCVP after excluding the families of ferns. Information on phylogenetic and distributional data availability taken from Rudbeck et al. (2022). Figure created using the “eulerr” package for R (Larsson, 2021).

## Notes

https://github.com/bmaitner/shortfalls

### Competing Interest Statement

The authors have declared no competing interest.

## References

Banki O, Hobern D, Döring M, Remsen D. 2019. Catalogue of Life Plus: A collaborative project to complete the checklist of the world’s species. Biodiversity Information Science and Standards; Sofia.

Bar-On YM, Phillips R, Milo R. 2018. The biomass distribution on Earth. Proceedings of the National Academy of Sciences of the United States of America 115: 6506–6511.

Boyle B, Hopkins N, Lu Z, Raygoza Garay JA, Mozzherin D, Rees T, Matasci N, Narro ML, Piel WH, McKay SJ, et al. 2013. The taxonomic name resolution service: an online tool for automated standardization of plant names. BMC bioinformatics 14: 16.

Boyle B, Maitner B. 2021. GNRS: Access the ‘Geographic Name Resolution Service’.

Brummitt RK, Pando F, Hollis S, Brummitt NA. 2001. World geographical scheme for recording plant distributions. York, UK: International Working Group on Taxonomic Databases for Plant Sciences.

Center for International Earth Science Information Network (CIESIN). 2018. Documentation for the Gridded Population of the World, Version 4 (GPWv4), Revision 11 Data Sets. Palisades NY: NASA Socioeconomic Data and Applications Center (SEDAC): 1–52.

Chamberlain S, Sagouis A. 2021. parzer: Parse Messy Geographic Coordinates.

Cheruvelil KS, Soranno PA. 2018. Data-Intensive Ecological Research Is Catalyzed by Open Science and Team Science. Bioscience 68: 813–822.

Cornwell WK, Pearse WD, Dalrymple RL, Zanne AE. 2019. What we (don’t) know about global plant diversity. Ecography 42: 1819–1831.

Díaz S, Kattge J, Cornelissen JHC, Wright IJ, Lavorel S, Dray S, Reu B, Kleyer M, Wirth C, Prentice IC, et al. 2016. The global spectrum of plant form and function. Nature 529: 167–171.

Enquist BJ, Condit R, Peet B, Schildhauer M, Thiers B, the BIEN working group. 2009. The Botanical Information and Ecology Network (BIEN): Cyberinfrastructure for an integrated botanical information network to investigate the ecological impacts of global climate change on plant biodiversity. The iPlant Collaborative.

Falster D, Gallagher R, Wenk EH, Wright IJ, Indiarto D, Andrew SC, Baxter C, Lawson J, Allen S, Fuchs A, et al. 2021. AusTraits, a curated plant trait database for the Australian flora. Scientific data 8: 254.

Feng X, Enquist BJ, Park DS, Boyle B, Breshears DD, Gallagher RV, Lien A, Newman EA, Burger JR, Maitner BS, et al. 2022. A review of the heterogeneous landscape of biodiversity databases: Opportunities and challenges for a synthesized biodiversity knowledge base. Global ecology and biogeography: a journal of macroecology 31: 1242–1260.

Fricke EC, Ordonez A, Rogers HS, Svenning J-C. 2022. The effects of defaunation on plants’ capacity to track climate change. Science 375: 210–214.

Gallagher RV, Allen S, Rivers MC, Allen AP, Butt N, Keith D, Auld TD, Enquist BJ, Wright IJ, Possingham HP, et al. 2020a. Global shortfalls in extinction risk assessments for endemic flora.

Gallagher RV, Falster DS, Maitner BS, Salguero-Gómez R, Vandvik V, Pearse WD, Schneider FD, Kattge J, Poelen JH, Madin JS, et al. 2020b. Open Science principles for accelerating trait-based science across the Tree of Life. Nature ecology & evolution 4: 294–303.

Geange SR, von Oppen J, Strydom T, Boakye M, Gauthier T-LJ, Gya R, Halbritter AH, Jessup LH, Middleton SL, Navarro J, et al. 2021. Next-generation field courses: Integrating Open Science and online learning. Ecology and evolution 11: 3577–3587.

Giuliani G, Peduzzi P. 2011. The PREVIEW Global Risk Data Platform: a geoportal to serve and share global data on risk to natural hazards. Natural Hazards and Earth System Sciences 11: 53–66.

Govaerts R, Nic Lughadha E, Black N, Turner R, Paton A. 2021. The World Checklist of Vascular Plants, a continuously updated resource for exploring global plant diversity. Scientific data 8: 215.

Guerrero-Ramírez NR, Mommer L, Freschet GT, Iversen CM, McCormack ML, Kattge J, Poorter H, Plas F, Bergmann J, Kuyper TW, et al. 2021. Global root traits (GRooT) database. Global ecology and biogeography: a journal of macroecology 30: 25–37.

Hortal J, de Bello F, Diniz-Filho JAF. 2015. Seven shortfalls that beset large-scale knowledge of biodiversity. Annual review of.

Institute for Economics and Peace. 2019. Global Peace Index 2019. Institute for Economics & Peace.

Iversen C, Powell A, McCormack M, Blackwood C, Freschet G, Kattge J, Roumet C, Stover D, Soudzilovskaia N, Valverde-Barrantes O. 2017. Fine-Root Ecology Database (FRED): A global collection of root trait data with coincident site, vegetation, edaphic, and climatic data, version 1.

Kattge J, Bönisch G, Díaz S, Lavorel S, Prentice IC, Leadley P, Tautenhahn S, Werner GDA, Aakala T, Abedi M, et al. 2020. TRY plant trait database - enhanced coverage and open access. Global change biology 26: 119–188.

Kattge J, Diaz S, Lavorel S, Prentice IC, Leadley P, Bönisch G, Garnier E, Westoby M, Reich PB, Wright IJ, et al. 2011. TRY--a global database of plant traits. Global change biology 17: 2905–2935.

Keller, Ankenbrand, Bruelheide, Dekeyzer, Enquist, Erfanian, Falster, Gallagher, Hammock, Kattge, et al. 2022. Ten (mostly) simple rules to future-proof trait data in ecological and evolutionary sciences. Authorea Preprints.

Larsson J. 2021. eulerr: Area-Proportional Euler and Venn Diagrams with Ellipses.

Lavorel S, Garnier E. 2002. Predicting changes in community composition and ecosystem functioning from plant traits: revisiting the Holy Grail. Functional ecology 16: 545–556.

LeBauer D, Kooper R, Mulrooney P, Rohde S, Wang D, Long SP, Dietze MC. 2018. BETYdb: a yield, trait, and ecosystem service database applied to second generation bioenergy feedstock production. GCB Bioenergy 10: 61–71.

Maitner B, Boyle B. 2022. TNRS: Taxonomic Name Resolution Service.

Maitner BS, Boyle B, Casler N, Condit R, Donoghue J, Durán SM, Guaderrama D, Hinchliff CE, Jørgensen PM, Kraft NJB, et al. 2017. The BIEN R package: A tool to access the Botanical Information and Ecology Network (BIEN) database. Methods in ecology and evolution /British Ecological Society 9: 373–379.

McGill BJ, Enquist BJ, Weiher E, Westoby M. 2006. Rebuilding community ecology from functional traits. Trends in ecology & evolution 21: 178–185.

Meijer JR, Huijbregts MAJ, Schotten KCG, Schipper AM. 2018. Global patterns of current and future road infrastructure. Environmental research letters: ERL [Web site] 13: 064006.

Migliavacca M, Musavi T, Mahecha MD, Nelson JA, Knauer J, Baldocchi DD, Perez-Priego O, Christiansen R, Peters J, Anderson K, et al. 2021. The three major axes of terrestrial ecosystem function. Nature 598: 468–472.

Mouillot D, Loiseau N, Grenié M, Algar AC, Allegra M, Cadotte MW, Casajus N, Denelle P, Guéguen M, Maire A, et al. 2021. The dimensionality and structure of species trait spaces. Ecology letters 24: 1988–2009.

Olson DM, Dinerstein E, Wikramanayake ED, Burgess ND, Powell GVN, Underwood EC, D’amico JA, Itoua I, Strand HE, Morrison JC, et al. 2001. Terrestrial Ecoregions of the World: A New Map of Life on Earth. BioScience 51: 933.

Penone C, Davidson AD, Shoemaker KT, Di Marco M, Rondinini C, Brooks TM, Young BE, Graham CH, Costa GC. 2014. Imputation of missing data in life-history trait datasets: which approach performs the best? Methods in Ecology and Evolution 5: 961–970.

R Core Team. 2020. R: A Language and Environment for Statistical Computing. Vienna, Austria: R Foundation for Statistical Computing.

Rousset F, Ferdy J-B. 2014. Testing environmental and genetic effects in the presence of spatial autocorrelation. Ecography 37: 781–790.

Rudbeck AV, Sun M, Tietje M, Gallagher RV, Govaerts R, Smith SA, Svenning J-C, Eiserhardt WL. 2022. The Darwinian shortfall in plants: phylogenetic knowledge is driven by range size. Ecography 2022.

Sandel B, Pavelka C, Hayashi T, Charles L, Funk J, Halliday FW, Kandlikar GS, Kleinhesselink AR, Kraft NJB, Larios L, et al. 2021. Predicting intraspecific trait variation among California’s grasses. The Journal of ecology 109: 2662–2677.

Sauquet H, von Balthazar M, Magallón S, Doyle JA, Endress PK, Bailes EJ, Barroso de Morais E, Bull-Hereñu K, Carrive L, Chartier M, et al. 2017. The ancestral flower of angiosperms and its early diversification. Nature communications 8: 16047.

Schneider FD, Fichtmueller D, Gossner MM, Güntsch A, Jochum M, König-Ries B, Le Provost G, Manning P, Ostrowski A, Penone C, et al. 2019. Towards an ecological trait data standard. Methods in ecology and evolution /British Ecological Society 10: 2006–2019.

Stein ML. 1999. Interpolation of Spatial Data: Some Theory for Kriging. Springer Science & Business Media.

Tavşanoğlu Ç, Pausas JG. 2018. A functional trait database for Mediterranean Basin plants. Scientific data 5: 180135.

Violle C, Navas M-L, Vile D, Kazakou E, Fortunel C, Hummel I, Garnier E. 2007. Let the concept of trait be functional! Oikos 116: 882–892.

Westoby M, Falster DS, Moles AT, Vesk PA, Wright IJ. 2002. Plant Ecological Strategies: Some Leading Dimensions of Variation between Species. Annual review of ecology and systematics 33: 125–159.

World Bank World Development Indicators. 2016. Government expenditure on education, total (% of government expenditure).

World Bank World Development Indicators. 2017. Research and development expenditure (% of GDP).

Wright IJ, Reich PB, Westoby M, Ackerly DD, Baruch Z, Bongers F, Cavender-Bares J, Chapin T, Cornelissen JHC, Diemer M, et al. 2004. The worldwide leaf economics spectrum. Nature 428: 821–827.

Zeballos SR, Giorgis MA, Cabido M, Gurvich DE. 2017. Unravelling the coordination between leaf and stem economics spectra through local and global scale approaches. Austral ecology 42: 394–403.

